# A Microengineered Brain-Chip to Model Neuroinflammation in Humans

**DOI:** 10.1101/2022.03.11.484005

**Authors:** I. Pediaditakis, K. R. Kodella, D. V. Manatakis, C. Y. Le, S. Barthakur, A. Sorets, A. Gravanis, L. Ewart, L. L. Rubin, E. S. Manolakos, C. D. Hinojosa, K. Karalis

**Author notes:** These authors contributed equally (I. Pediaditakis, K. R. Kodella).

## Abstract

Species differences in the brain and the blood-brain barrier (BBB) biology hamper the translation from animal models to humans and impede the development of specific therapeutics for brain diseases. Here we present a human Brain-Chip engineered to recapitulate critical aspects of the complex brain cell-cell interactions that mediate neuroinflammation development. Our human organotypic microphysiological system (MPS) includes endothelial-like cells, pericytes, glia, and cortical neurons and maintains BBB permeability at *in vivo* relevant levels, providing a significant improvement in complexity and clinical mimicry compared to previous MPS models. This is the first report of a Brain-Chip with an RNA expression profile close to that of the adult human cortex and that demonstrates advantages over Transwell culture. Through perfusion of TNF-α, we recreated key inflammatory features, such as glia activation, the release of proinflammatory cytokines, and increased barrier permeability. Our model may provide a reliable tool for mechanistic studies in neuron-glial interactions and dysregulation of BBB function during neuroinflammation.

## INTRODUCTION

Neurodegenerative diseases (ND) are a serious public health problem, with their increasing burden accounting for more than one billion affected people worldwide (Carroll, 2019). Although the molecular mechanisms underlying changes in brain cell-to-cell interactions associated with ND have been elucidated to a significant extent, therapeutic targets and predictive biomarkers are still lacking. Emerging evidence points to inflammation as a major pathogenetic mechanism for neuropathology underlying neurodegenerative diseases (Guzman-Martinez et al., 2019a; Minghetti, 2005). Growing evidence indicates that brain events, such as local ischemia and systemic inflammatory conditions, create a microenvironment favoring the entry of peripheral activated immune cells, or their secreted proinflammatory factors, into the brain by compromising the integrity of the blood-brain barrier (BBB) (Perry et al., 2007a). There is a large body of evidence for a pivotal role of microglia and astrocytes in the pathogenesis of neurodegenerative disorders (Crotti and Ransohoff, 2016; Kim and Joh, 2006; Liddelow and Barres, 2017; Liddelow et al., 2017; Linnerbauer et al., 2020; Perry et al., 2010). Activation of microglia is characterized by the production of cytotoxic molecules and pro-inflammatory cytokines, affecting cellular homeostasis, and ultimately establishing a microenvironment driving neuronal damage (Perry et al., 2010).

In addition, reactive astrocytes are also found to be implicated in impaired neuronal function and survival promoting inflammation and increasing cell damage in the CNS (Horng et al., 2017; Liddelow and Barres, 2017; Linnerbauer et al., 2020; Sofroniew, 2020). Additional reports suggest a crosstalk between astrocytes and microglia in the context of neuroinflammation and neurodegeneration (Linnerbauer et al., 2020). Reactive microglia and astrocytes are likely to contribute to the leaky BBB observed in these diseases through downregulation of paracellular tight-junction proteins such as Occludin and zonula occludens-1(ZO-1) (Keaney and Campbell, 2015; Obermeier et al., 2013).

Despite the progress in understanding the mechanisms mediating the effects of immune activation in the brain (González et al., 2014), the translation of these findings to effective, specific treatments lags significantly. Species differences between human and the currently used experimental *in vivo* animal models of ND, together with the inherent limitations of the primary brain rodent cells or human cell lines currently used as *in vitro* models, are greatly implicated in this problem. Advances in stem cell biology have recently enabled new human cell models for experimentation, such as iPSC-derived brain cells and complex, multi-layered 3D organoids (Bose et al., 2021; Dolmetsch and Geschwind, 2011; Yin et al., 2016). These systems can provide new insights into tissue biology and our understanding of the interindividual differences in brain functions. Exploiting these systems can be significantly advanced by their culture in microphysiological platforms providing a perfusable vascular system and an overall physiologically relevant microenvironment, by enabling long-lasting cell interactions and the associated functionality (Cucullo et al., 2011). Recent reports describe applications of this technology in developing Organ-Chips that recapitulate aspects of the complex BBB functions (Ahn et al., 2020; Maoz et al., 2018; Park et al., 2019; Vatine et al., 2019). These microfluidic models have certainly overcome several of the limitations of traditional culture models. However, so far, they have not included microglial cells, an essential component of the neurovascular unit (NVU), critically involved in the regulation of neuroinflammatory activity (McConnell et al., 2017; Muoio et al., 2014; Streit and Kincaid-Colton, 1995).

In pathological states, microglia and astrocytes undergo complex changes increasing their capacity to produce proinflammatory cytokines such as tumour necrosis factor-alpha (TNF-α) (Lau and Yu, 2001; Wang et al., 2015), interleukin-1β (IL-1β), interleukin-6 (IL-6), and interferon-gamma (IFN-γ), that increase the BBB permeability (De Vries et al., 1996; Yarlagadda et al., 2009). TNF-α was shown to increase BBB permeability, induce the activation of glia *in vivo* (Cheng et al., 2018; Neniskyte et al., 2014a), and potentiate glutamate-mediated cytotoxicity, a process linked to neuronal death (Olmos and Lladó, 2014). Elevated levels of TNF-α have been found in traumatic brain injury (Frugier et al., 2010), ischemic stroke (Zaremba and Losy, 2001), Alzheimer’s (AD) (Jiang et al., 2011), Parkinson’s (PD) (Kouchaki et al., 2018), Multiple Sclerosis (MS) (Rossi et al., 2014), and amyotrophic lateral sclerosis (ALS) (Cereda et al., 2008).

Here we describe how we leveraged the Human Emulation System^®^ to build a comprehensive Brain-Chip model to characterize cellular interactions underlying the development of neuroinflammation. We have populated our Brain-Chip with human primary astrocytes, pericytes, and iPSC-derived brain microvascular endothelial cells, together with human iPSC-derived cortical neurons and microglial cell line. This Brain-Chip supported a tissue relevant, multicellular architecture and the development of a tight blood brain barrier, sustained over seven days in culture. Using next-generation sequencing data and information retrieved from well-curated databases of signature gene sets for the human cortex, we demonstrate that the Brain-Chip’s transcriptomic signature is closer to the adult cortical tissue than the conventional cell culture systems used to study the BBB *in vitro* (Stone et al., 2019; Thomsen et al., 2015).

To simulate changes occurring during the early phases of neuroinflammation, we perfused either the brain or the vascular side of the Brain-Chip with TNF-α. We successfully reproduced key clinical features of neuroinflammation, such as disruption of the BBB, astrocyte, and microglial activation, increased proinflammatory cytokine release, capturing contributions of the individual brain cell types to inflammatory stimuli. In summary, we present an *in vivo* relevant human Brain-Chip as a multicellular model designed to investigate neurodegenerative pathogenesis and future applications, including clinical studies that could lead to effective precision medicine treatments.

## RESULTS

### Microengineered Human Brain-Chip Platform of the Neurovascular Unit

We leveraged organ-on-chip technology and the recent progress in developing human brain primary and iPSC-derived differentiated cells to generate a human Brain-Chip model that enables the stimulation and monitoring of inflammatory responses. As described previously, the Brain-Chip has two microfluidic channels, separated by a thin, porous polydimethylsiloxane (PDMS) membrane that enables cellular communication and supports coating with tissue-specific extracellular matrix (ECM) (Pediaditakis et al., 2021). The top channel, we refer to as the “brain” channel of the chip, accommodates the co-culture of key elements of the neurovascular unit (NVU), including excitatory and inhibitory cortical neurons, microglia, astrocytes, and pericytes (McConnell et al., 2017). The bottom channel, which we refer to as the “vascular” channel of the chip, is seeded with human iPSC-derived brain microvascular endothelial-like cells (iBMECs) that create a lumen-like structure (Wong et al., 2013), modelling the interface between the circulation and the brain parenchyma (**Figure 1A**). We used cell proportions comparable to those reported (von Bartheld et al., 2016; Shepro and Morel, 1993; Sultan and Shi, 2018; Xu et al., 2010), with the understanding of the challenge in recapitulating the *in vivo* cellular milieu.

**Figure 1.**
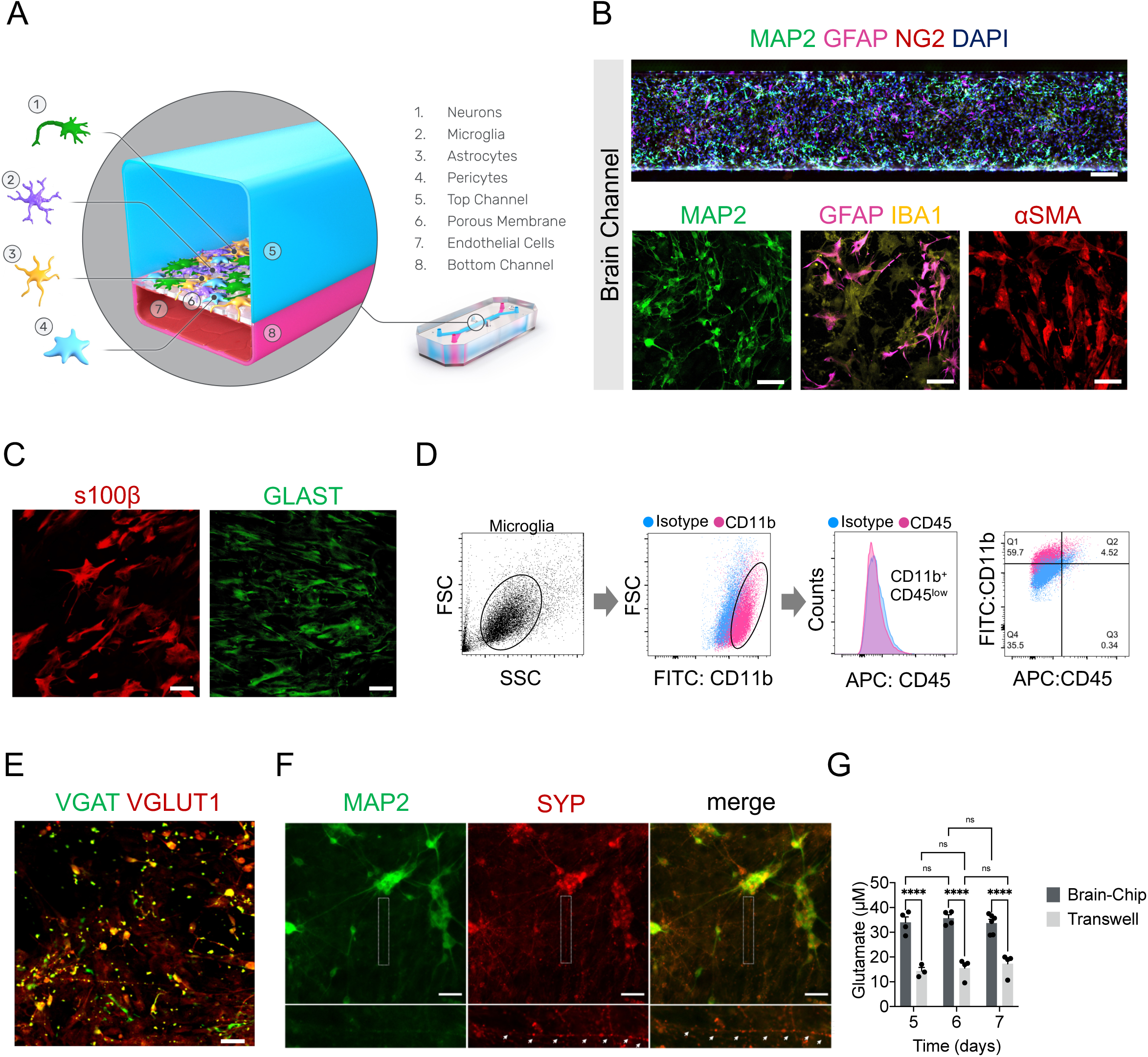
Reconstruction of the neurovascular unit in the Brain-Chip. **(A)** Schematic illustration of the Brain-Chip, a two-channel micro-engineered chip including iPSC-derived brain endothelial-like cells cultured on all surfaces of the bottom channel, and iPSC-derived Glutamatergic and GABAergic neurons, primary human brain astrocytes, pericytes, and microglia on the surface of the top channel. **(B)** Confocal images of the cell coverage in the brain channel on day 7 of culture. Top image: Immunofluorescence staining of the brain channel including MAP2 (green), GFAP (magenta), NG2 (red), and DAPI (blue). Bottom images: Representative merged confocal image of the brain channel on culture day 7, stained for neurons (MAP2, green), astrocytes (GFAP, magenta, IBA1, yellow), and pericytes (αSMA, red) (bar, 50 μm). **(C)** Representative immunofluorescent staining for s100β (red) and GLAST (green) (bar,100 μm). **(D)** FACS analysis of cell specific markers of microglia: Total population of microglia within the brain channel (grey), CD11b positive population (magenta), CD45 positive population (magenta), quantification of CD11b:CD45 positive cells. **(E)** Representative merged confocal image of the brain channel co-stained with VGAT (green) for GABAergic neurons and VGLUT1 (red) for Glutamatergic neurons (bar, 100 μm). **(F)** Immunofluorescence staining of the brain channel including MAP2 (green), and SYP (red) (bar, 100 μm). **(G)** Levels of secreted glutamate in the brain channel on culture days 5, 6 and 7 (n=4-6 independent chips, ****P<0.0001, NS: not significant compared to the transwells group n=3-4). Data are expressed as mean ± S.E.M, statistical analysis by two-way ANOVA with Tukey’s multiple comparisons test.

To characterize the multicellular structure in the brain channel of the chip, we used specific markers for each brain cell type, including microtubule-associated protein 2 (MAP2) for neurons, glial fibrillary acidic protein (GFAP), s100β, and glutamate transporter (GLAST) for astrocytes, ionized calcium-binding adaptor protein-1 (IBA-1) for CD11b^+^ CD45^low^ microglia, neuron-glial antigen 2 (NG2), and smooth muscle alpha-actin (αSMA) for pericytes (**Figures 1B, 1C and 1D**). Positive staining for the vesicular glutamate transporter (VGLUT1) and the vesicular GABA transporter (VGAT) (**Figure 1E**), indicate the co-existence of excitatory and inhibitory neurons, respectively. To get an initial evaluation of the functionality of the neurons in the Brain-Chip, we performed immunofluorescent staining with synaptophysin (SYP), a nerve terminal marker synaptic vesicle that stains mature presynaptic neurons. Our results show that MAP2 positive cells expressed the synaptic marker synaptophysin (**Figure 1F**). The efficiency of synaptic transmission is governed by the probability of neurotransmitter release, the amount of neurotransmitter released from the presynaptic terminal. Additionally, our data demonstrate the functional maturation of the human iPSC-derived neurons in the Brain-Chip compared to transwells, as depicted by the secreted glutamate levels (**Figure 1G**), suggesting functional differences between the two systems. To the best of our knowledge, this is the first report on a human microphysiological system where inhibitory and excitatory cortical neurons and primary microglia complement the cellular components of BBB to form the neurovascular unit.

The establishment of the endothelial monolayer was assessed via staining for the tight junction-specific marker, zona occludens-1 (ZO-1), Occludin, as well as the endothelial-specific junctions PECAM1, which demonstrated that the endothelial-like cells form a consistent morphology along the entire vascular channel of the chip (**Figure 2A**). Additionally, we screened the endothelial-like cells we employed in our model for expression of the epithelium markers Cadherin1 (CADH1), TRPV6, and Claudin-4. As shown, we confirmed lack of expression in the endothelial-like cells, in contrast to the human epithelial cell lines used as positive controls (**Figure S1A)**. Once the endothelial-like cells are cultured juxtaposed to pericytes, astrocytes, microglia, and neurons in the Brain-Chip, they establish a tight barrier for seven days (**Figure 2B**). To this end, we evaluated the apparent permeability (Papp) in Brain-Chips seeded with human iPSC-derived brain microvascular endothelial-like cells from two different healthy donors (Donor 1; RUCDR, Donor 2; iXcell) (**Figure 2B**). The obtained Papp values in our model to 3 kDa, 10 kDa, 40 kDa, and 70 kDa dextran reached values as low as those reported from *in vivo* studies (Shi et al., 2014; Yuan et al., 2009) (**Figure 2C**), routinely cited as golden standards of apparent permeability. Moreover, the permeability values obtained in the Brain-Chip were comparable within a specified range to those reported by previous BBB studies with organ chips (Ahn et al., 2020; Vatine et al., 2019).

**Figure 2.**
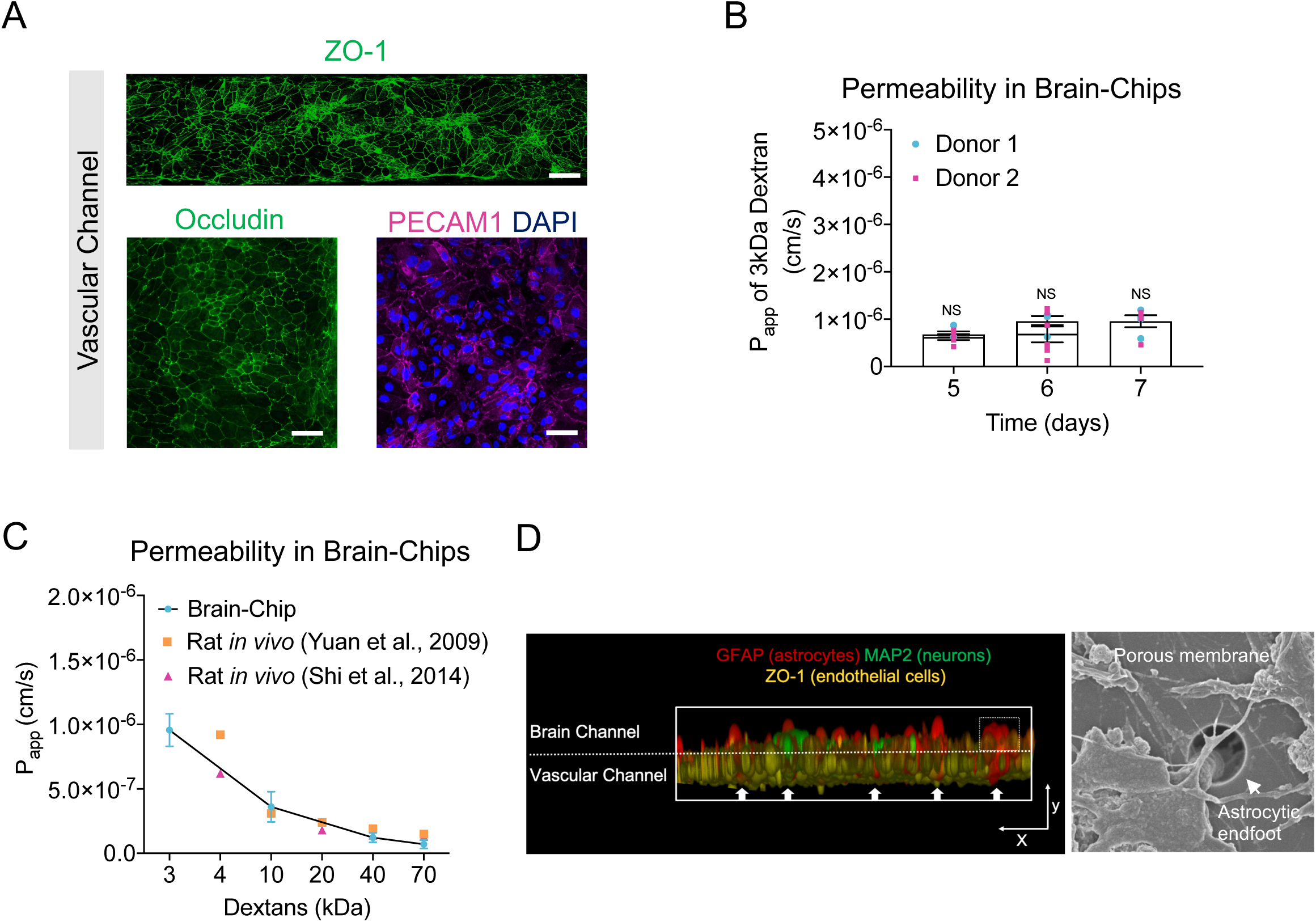
Characterization of the Barrier in the Brain-Chip. **(A)** Top Image: Immunofluorescence staining of the vascular channel stained for the tight junction marker ZO-1 (green) (bar, 1 mm). Bottom Images: Immunofluorescence micrographs of the human brain endothelium cultured on-chip for 7 days labeled with Occludin (green), PECAM1 (magenta), and DAPI (blue) (bar, 100 μm). **(B)** Quantitative barrier function analysis by the apparent permeability to 3kDa fluorescent dextran, in two independent iPSC donor lines on culture days 5, 6, and 7 (n=4-6 independent chips). NS: not significant. Data are expressed as mean ± S.E.M., statistical analysis by Student’s t-test. **(C)** The apparent permeability of different size dextran molecules (3-70 kDa) across Brain-Chips correlated with previously reported *in vivo* rodent brain uptake data. (n=3-5 independent chips). Data are expressed as mean ± S.E.M.) **(D)** Left: Exploded view of the chip. Interaction of primary human astrocyte end-feet-like processes (GFAP, red) with endothelial-like cells (ZO-1, yellow), MAP2 (green). Right: Representative Scanning Electron Microscopy (SEM) image showing astrocytic endfoot passing through 7 µm pores into the vascular channel (bar, 30um). White arrows show the astrocytic endfoot.

Findings on microglial location in the perivascular space highlight their interaction with endothelial cells and support their influence on BBB integrity although, very few studies have been conducted to delineate a direct link between microglia and barrier function (Haruwaka et al., 2019a). On the other hand, astrocytes by providing a connection between the endothelial blood flux and neurons (**Figure 2D**), are critical for the formation and maintenance of the BBB (Alvarez et al., 2013).To explore the potential impact of microglia and astrocytes on the barrier integrity, we measured the permeability to cascade-blue 3-kDa dextran. Significant decrease in paracellular permeability in the presence of microglia highlights its importance in the maintenance of the barrier integrity in the Brain-Chip (**Figure S1B**). Further, we found significant increase in the permeability in the absence of astrocytes, in line with their reported significant role in the stability and maintenance of the BBB (Abbott et al., 2006). No significant effect in the permeability was detected by eliminating neurons, the most abundant cell in the Brain-Chip. Taken together, all the above support the hypothesis that the decrease in permeability in the Brain-Chip following challenge with TNF-α is specifically driven from the microglia and the astrocytes, rather than from a technical/mechanical obstacle, such as clogging of the pores.

Furthermore, the comparison to chips that contain only iPSC-derived brain microvascular endothelial-like cells provides further reassurance on the contribution of the supporting cells in the Brain-Chip that also contains pericytes, astrocytes, microglia, and neurons (Keaney and Campbell, 2015; Obermeier et al., 2013) (**Figure S1C**) in the stability of the barrier. We also confirmed that the cells in the vascular channel expressed the brain endothelium-specific glucose transporter, GLUT-1 (Veys et al., 2020) (**Figure S1D**), and showed internalization of transferrin (**Figure S1E**), an essential mechanism for transport across the BBB leveraged for delivery of therapeutic antibodies (Jones and Shusta, 2007). Further characterization of iBMECs via FACS analysis confirmed the expression of the brain endothelium-specific glucose transporter, GLUT-1, transferrin receptor, and efflux transporters in the BBB, such as P-glycoprotein (P-gp) and MRP-1 (**Figure S1F**), of additional reassurance on the potential of the model in evaluating the ability of developing therapeutics to enter the brain.

### Transcriptomic Comparison of the Human Brain-Chip Versus Conventional Transwell Systems and Adult Human Tissue

After confirming the *in vivo* relevant cell composition and barrier function, we assessed the extent of similarities in gene expression between the Brain-Chip and the adult human cortex tissue, as well as the differences compared to the transwell brain model, the most commonly used cell culture system for modeling brain *in vitro*. Using the same cell composition and experimental conditions in transwells and Brain-Chip cultures, we performed RNAseq analyses on days 5 and 7 of culture. Expression of specific markers for each of the cell types seeded, confirmed their representation in the culture at the time of the analyses. Principal Component Analysis (PCA) showed clear separation between the samples of the two models in the 2-dimensional space determined by the first two PCs explaining 47.5% of the total variance in the data (**Figure 3A**). Unlike transwells, Brain-Chips were clustered together in the 2D PCA space, which indicates their transcriptomic “stability” across the days in culture (**Figure 3A**). Next, we examined the differential gene expression (DGE) in Brain-Chips compared to transwells. Out of the 57,500 genes annotated in the genome, 5695 were significantly differentially expressed (DE) between these samples: 3256 and 2439 genes were up- and down-regulated, respectively (**Figure 3B**). Next, using the information of the up- and down-DE genes, we performed gene ontology (GO) enrichment analysis to identify the significantly enriched biological processes in the two systems (Ashburner et al., 2000; Mi et al., 2013). In Brain-Chips we identified significantly enriched pathways (FDR p-value ≤ 0.05) of the Brain channel related to the extracellular matrix organization, cell adhesion, and tissue development (**Figure 3C**). Evidence has been accumulated that activity-dependent aggregation and proteolysis of ECM (extracellular matrix organization) and associated molecules shape synaptogenesis, synapse maturation, and synaptic circuit remodeling (Ferrer-Ferrer and Dityatev, 2018). Extracellular matrix accelerates the formation of neural networks and communities in a neuron-glia co-culture (Lam et al., 2019). In contrast, axongenesis, axon guidance, chemotaxis, neurogenesis, neuron migration, and cell differentiation pathways were significantly enriched in transwells (**Figure 3D**), supporting the hypothesis of incomplete neuronal maturation in these systems (Hesari et al., 2016; Sances et al., 2018). A basic property of immature neurons is their ability to change position from the place of their final mitotic division in proliferative centers of the developing brain to the specific positions they will occupy in a given structure of the adult nervous system (Rakic, 1990). Proper acquisition of neuron position, attained through the process of active migration and chemotaxis (Cooper, 2013).

**Figure 3.**
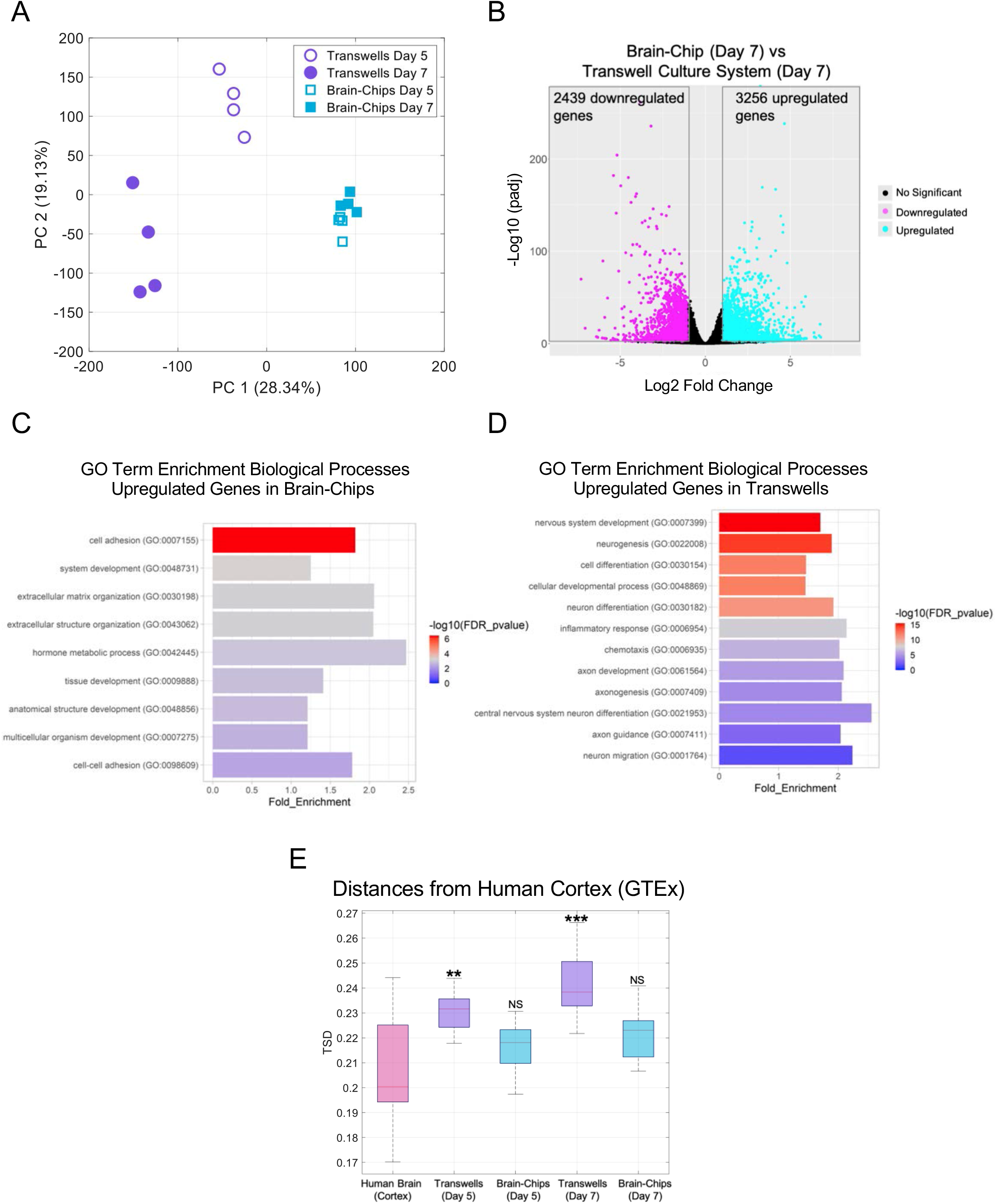
Comparative analyses of the transcriptomic profiles of the Brain-Chip, adult cortex tissue and transwell culture. **(A)** Principal Component Analysis (PCA) of RNAseq data from the brain channel of the Brain-Chip and the transwell brain cells culture on culture days 5 and 7 (n=4 per condition). The first two PCs explain the 47.47% of the total variance. **(B)** DGE analysis Identified up (cyan) - and down (magenta) - regulated genes (dots) in the Brain-Chip as compared to transwells on culture day 7. (**C** to **D**) Biological processes in Brain-Chip and transwells, as identified by Gene Ontology (GO) enrichment analysis based on the DE genes. (**E**) Boxplots summarizing the distributions of the corresponding pairwise TSD distances. In each pair, one sample belongs to the reference tissue (Human Brain-Cortex) and the other either to the reference tissue or to one of our culture models, i.e., Brain-Chip or transwell, from culture days 5 and 7. The Brain-Chip and transwell cultures were run in parallel. n=4 independent chips; data are expressed as mean ± S.E.M. NS: Not Significant, **P<0.01, ***P<0.001; statistical analysis with two-sample t-test using a null hypothesis that the data from human tissue and the data from chips or transwells comes from independent random samples from normal distributions with equal means and equal but unknown variances. On each box the red line indicates the median and the bottom and top edges correspond to the 25^th^ and 75^th^ percentiles respectively. The whiskers extend to the most extreme but not considered outliers values.

Lastly, we used the Transcriptomic Signature Distance (TSD) (Manatakis et al., 2020) to calculate the Brain-Chip’s and transwell’s transcriptomic distances from the adult human cortex tissue. We were able to show that the Brain-Chip exhibits higher transcriptomic similarity (smaller transcriptomic distance) to the adult human cortex on either day of culture compared to the transwells (**Figure 3E**), by leveraging next-generation sequencing data and information retrieved from the Human Protein Atlas database providing signature gene sets characteristic for the human brain (Uhlén et al., 2015). To further support our conclusions, we used the cerebral cortex RNA-seq data, from two different human donors (donor 9861 and 10021), available in the Allen Brain Atlas (Available from: human.brain-map.org). For each donor, we measured the transcriptomic signature distances (TSDs) between the corresponding cerebral cortex samples and (i) Brain-Chip and (ii) Transwells on Days 5 and 7. For both donors, the results clearly indicate that for both days, our Brain-Chips are statistically significantly closer to human cerebral cortex as compared to transwells (**Figure S2**). Cumulatively, these findings demonstrate that the Brain-Chip recapitulates the cortical brain tissue more closely than conventional cell culture systems such as transwells.

### TNF-α-Induced Neuroinflammation in the Brain-Chip

Neuroinflammation is emerging as a key mechanism in the progress of several infectious and neurodegenerative diseases (Guzman-Martinez et al., 2019b; Perry et al., 2007a). While the pathways driving neuroinflammation in response to infection have been delineated, the exact mechanisms that trigger and sustain the activation of glia (microglia and astrocytes) and the direct association with the barrier function remain unclear.

To model neuroinflammation in the Brain-Chip, we perfused TNF-α directly in the brain channel at a concentration of 100 ng/mL starting on day five of culture (**Figures 4A and S3A**) for two days. The concentration of 100 ng/mL TNF-α was chosen based on previous studies, where this concentration stimulated proinflammatory factors to be released and compromised the barrier integrity (Rochfort et al., 2014, 2016). As the majority of neuroinflammatory responses are elicited by the supportive glial cells, in particular microglia and astrocytes, the incorporation and evaluation of these cell types in *in vitro* assay systems is of particular interest in the pharmaceutical field.

**Figure 4.**
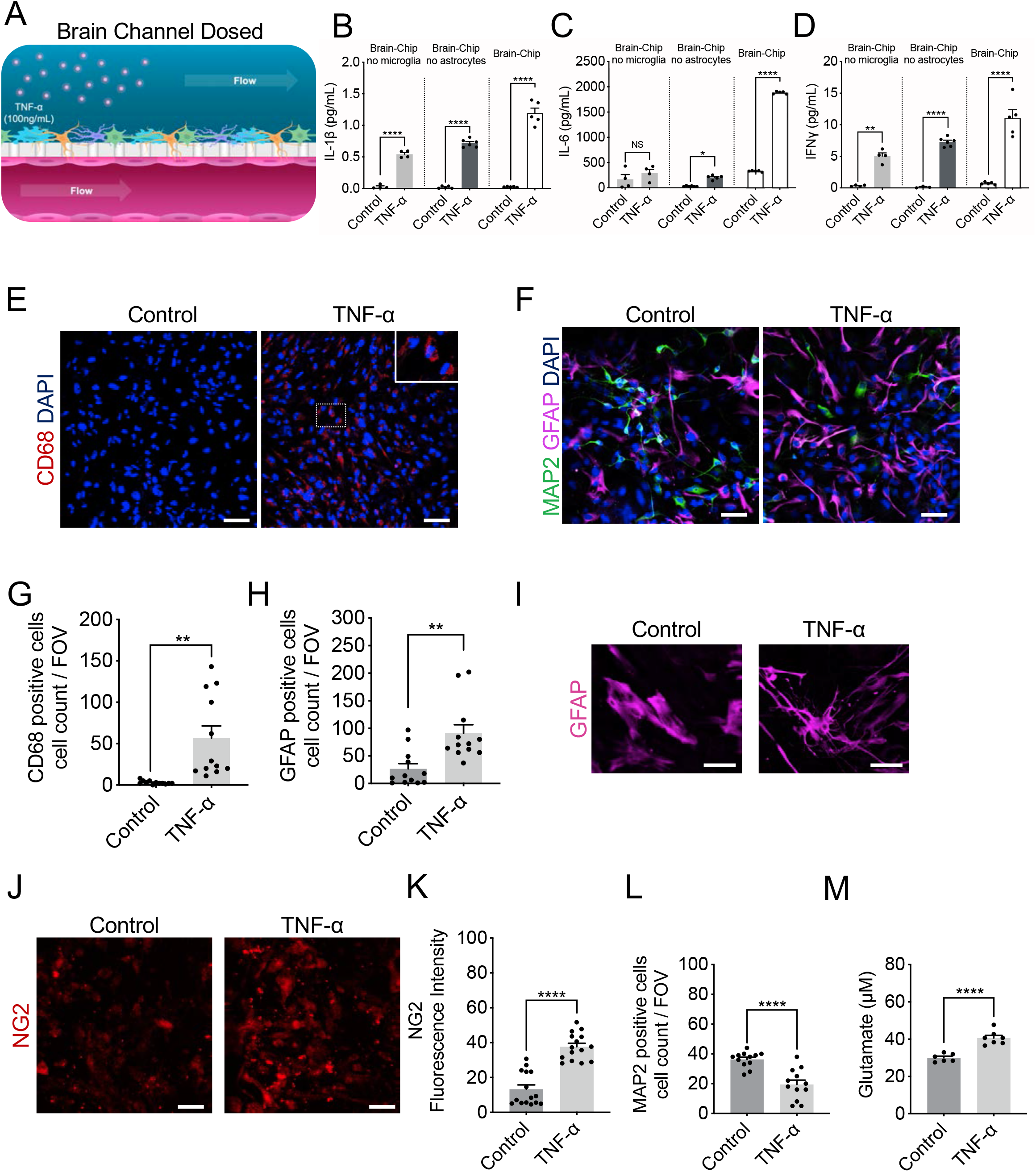
Response of the Brain-Chip to neuroinflammation. (**A**) Schematic illustration of the induction of neuroinflammation by perfusion of TNF-α through the brain channel. (**B** to **D**) Secreted levels of IL-1β, IL-6, and IFNγ in control or TNF-α-treated Brain-Chips including, or not, microglia and/or astrocytes. n=4-5 independent chips; data are expressed as mean ± S.E.M. NS: Not Significant, *P<0.05, **P<0.01, ****P<0.0001; statistical analysis with Student’s t-test. (**E** to **F**) Representative immunofluorescent staining for microglia (CD68, red), neurons (MAP2, green), astrocytes (GFAP, magenta), and nuclei (DAPI, blue) in TNF-α-treated of control chips (bar, 100 nm). **(G)** Quantification of the CD68-positive events/field of view in 4 randomly selected different areas/chip, n=3 Brain-Chips; data are expressed as mean ± S.E.M., \*\**P*<0.01 compared to the untreated control group, statistical analysis by Student’s t-test. **(H)** Quantification of the number of GFAP-positive and MAP2 events/field of view in n=4 randomly selected different areas/chip, n=3 Brain-Chips, \*\**P*<0.01, compared to the untreated control group; statistical analysis with Student’s t-test. **(I)** Immunofluorescence images show an example of the two types of astrocyte morphology in cultures (polygonal shape towards more elongated shape), after immunostaining with an antibody against GFAP (bar, 100 nm). **(J)** Representative immunofluorescent staining for pericytes (NG2, red) in TNF-α-treated or control chips (bar, 100 nm). **(K)** Quantification of NG2 fluorescent intensity in n=5 randomly selected different areas/chip, n=3 Brain-Chips; data are expressed as mean ± S.E.M., \*\*\*\**P*<0.0001 compared to the untreated control group; statistical analysis with Student’s t-test. **(L)** Quantification of the number of MAP2 events/field of view in n=4 randomly selected different areas/chip, n=3 Brain-Chips, \*\*\*\**P*<0.0001, compared to the untreated control group; statistical analysis with Student’s t-test. **(M)** Levels of secreted glutamate in the brain channel on culture day 7 (n=6-7 independent chips. \*\*\*\**P*<0.0001, compared to the untreated group. Data are expressed as mean ± S.E.M, statistical analysis by Student’s t-test.

To confirm the advantage of the Brain-Chips containing both microglia and astrocytes, the key cellular mediators of neuroinflammatory processes, we compared TNF-α-induced secretion of inflammatory cytokines in Brain-Chip in the presence or absence of microglia or astrocytes in the brain channel. We found significantly increased levels of interleukin-1β (IL-1β), interleukin-6 (IL-6), and interferon-gamma (IFNγ) in the effluent of the Brain-Chips containing microglia and astrocytes, following 48 h of exposure to TNF-α (**Figures 4B to D)**. Notably, absence of microglia resulted in compromised induction of both IL-1β and IFNγ, while it prevented any change in IL-6 secretion, in contrast to the high levels of IL-6 detected in the microglia-containing Brain-Chip (**Figures 4B to D)**. These findings capture the microglia-dependent inflammatory responses to TNF-α in the Brain-Chip, in line with previous studies.

These findings demonstrate that Brain-Chips containing microglia respond to noxious stimuli such as TNF-α by mounting an inflammatory response in a manner similar to that shown in *in vivo* studies (Block et al., 2007; Brás et al., 2020; Haruwaka et al., 2019b; Kuno et al., 2005; Merlini et al., 2021; Perry et al., 2007b).

Although microglia cells are the primary source of cytokines, astrocytes do also perpetuate the destructive environment via secretion of various chemokines and proinflammatory cytokines, including IL-1β and IFNγ (Rothhammer and Quintana, 2015), in line with our findings. Interestingly, TNF-α induced IL-6 secretion directly from the microglia but not from the astrocytes, that required the presence of microglia for induction of cytokines secretion. Cumulatively, our data suggest the operation of different regulatory mechanisms and cell-to-cell interactions driving the cytokine responses of glial cells during inflammation. These findings indicate the potential of the Brain-Chip to critically support the development of specific new therapeutic approaches for patients.

In line with the increased cytokine release, TNF-α exposure led to activation of microglia and astrocytes, as depicted by significant increases in CD68 positive cells or the proportion of GFAP-positive cells respectively (**Figures 4E to 4H and S3B, S3C**), recapitulating *in vivo* findings for glial cells that constitute the first response to inflammatory and infectious stimuli (Boche et al., 2019; Sun et al., 2008). Astrocytes exposed to TNF-α displayed a significant change in their morphology, transitioning from a polygonal shape towards a more elongated (stellate) shape (**Figure 4I**). Further, we detected pericyte activation, in line with the notion of the importance of NG-2 reactive pericytes in neuroinflammation (Ferrara et al., 2016) (**Figure 4J and 4K**), although we did not detect any significant effect on the total nuclei count (**Figure S3D**). Our studies revealed no differences in the abundance of Ki67-positive cells between the control and treated groups, suggestive of TNF-α-induced activation of the existing, astrocytes, rather than generation of new astrocytes (**Figures 3SE and 3SF**).

In addition, exposure to TNF-α resulted in the loss of neuronal immunoreactivity of the cytoskeletal microtubule-associated protein (MAP-2) compared to the control, in neurons, suggesting that TNF-α induced neuronal injury/damage (**Figures 4F and 4L**), in line with the described neurotoxic effects of inflammatory mediators (Neniskyte et al., 2014b). Excessive brain TNF-α levels have been associated with the compromised activity of the glutamate transporters, which results in an increase in glutamate levels. The secreted glutamate levels in the effluent of the brain channel of the chip in control Brain-Chips remained stable on days 5, 6 and 7 of culture **(Figure 1G)**, while they increased significantly increased following two days of exposure to TNF-α (**Figure 4M)**. These data corroborate reports from *in vitro* and *in vivo* studies linking glutamate-induced excitotoxicity to neuroinflammation (Olmos and Lladó, 2014; Rossi et al., 2014; Ye et al., 2013).

Strong experimental evidence has demonstrated the multifaceted effects of TNF-α on the BBB anatomy and function, via its direct action on the endothelium as well as the downstream effects via induction of associated proinflammatory factors (Trickler et al., 2005; Zhao et al., 2007). We found that in the Brain-Chip, the perfusion of the brain channel with TNF-α (100 ng/mL) for two days, ensued loss of the integrity of the tight junctions, as demonstrated by diffuse ZO-1 staining (**Figures 5A**). Furthermore, the expression of intercellular adhesion molecule 1 (ICAM-1), a hallmark of inflammation-promoting adhesion and transmigration of circulating leukocytes across the BBB (Marchetti and Engelhardt, 2020), was also significantly induced (**Figure 5A**). Assessment of the barrier function revealed a significant increase in permeability to 3kDa dextran in the TNF-α treated Brain-Chip, in a time dependent-manner (**Figure 5B**). Furthermore, we show that the changes in barrier permeability were only evident in the presence of microglia (**Figure 5C**), confirming previous findings on the critical role of this cell in driving BBB dysfunction (Nishioku et al., 2010). To further support this hypothesis, we used Minocycline to inhibit the induction of reactive microglia (Ai et al., 2005; Henry et al., 2008). Reportedly, Minocycline reduces the characteristic BBB leakage in rodent models of brain diseases, such as hypoxia, ischemia, and Alzheimer’s disease (Ryu and McLarnon, 2006; Yang et al., 2015; Yenari et al., 2006). Our findings add on our understanding on barrier function in the human Brain-Chip as a result of TNF-α induced inflammation, as they also include the response of microglia (**Figure 5D**).

**Figure 5.**
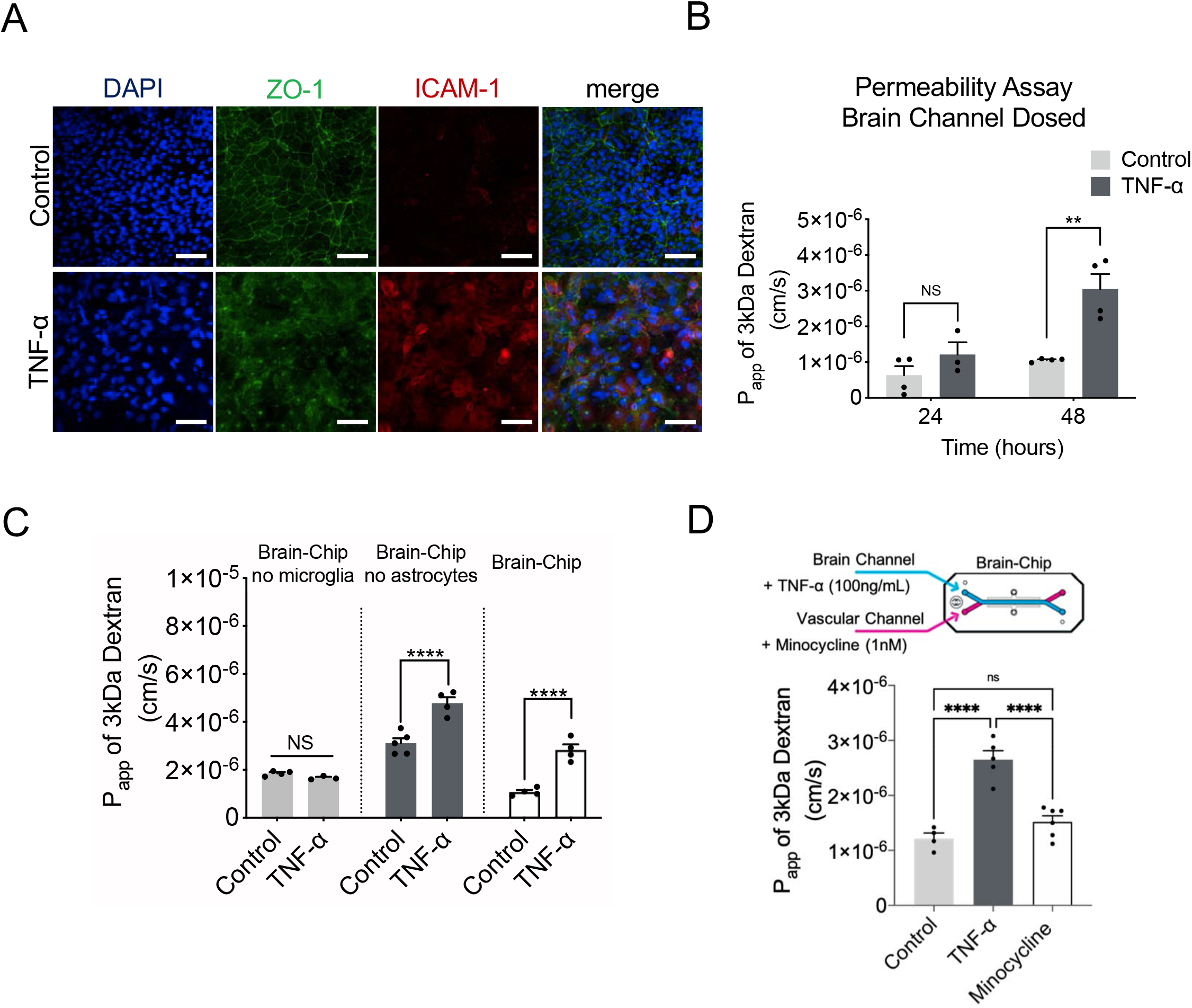
Barrier changes during neuroinflammation. **(A)** Representative merged image of immunofluorescent staining of Intercellular Adhesion Molecule 1 (ICAM-1, red), tight junction protein 1 (ZO-1, green), and cell nuclei (DAPI, blue), (bar, 100 nm). **(B)** Quantification of barrier permeability to 3 kDa fluorescent dextran, upon 24 and 48 h of treatment with TNF-α; n=3-4 independent chips. Data are expressed as mean ± S.E.M, NS: Not Significant, \*\**P*<0.01, control compared to TNF-α treated group; statistical analysis by two-way ANOVA followed by Tukey’s multiple comparisons test. **(C)** Assessment of the permeability of the Brain-Chip on culture day 7, in the absence or presence of microglia, astrocytes or neurons; n=4-8 independent chips; data are expressed as mean ± S.E.M., ****P<0.0001, NS: Not Significant compared to full model (Brain-Chip), statistical analysis by one-way ANOVA with Sidak’s multiple comparisons test. **(D)** Quantification of barrier apparent permeability to 3 kDa fluorescent dextran, upon 48 h of TNF-α-treated Brain-Chips including minocycline, or not (control); n=3-5 independent chips. Data are expressed as mean ± S.E.M, \*\*\*\**P*<0.0001, NS: Not Significant compared to TNF-α treated group or minocycline group; statistical analysis by Student’s t-test.

These results taken together show how the Brain-Chip can be applied to characterize specific changes in cell-cell interactions underlying the development and progress of neuroinflammation.

### Neuroinflammation in the Brain-Chip Induced by Vascular Exposure to TNF-α

Although pathology locally in the brain is associated with massive production of proinflammatory cytokines (neuroinflammation), reports show that systemic infections/inflammatory states spreading to the brain through the vascular system, can also induce inflammation in the brain and alter the progression of chronic neurodegenerative diseases (Perry, 2004). To assess how systemic (through the vascular channel) administration of TNF-α affects the cells in the brain channel (**Figure 6A and S4A**), we first performed immunostaining for CD68, GFAP, and NG2 two days after vascular administration of TNF-α, that revealed activation of microglia, astrocytes, and respectively (**Figure 6B to 6F**), while we did not detect any significant effect on the total nuclei count (**Figure 6G**).

**Figure 6.**
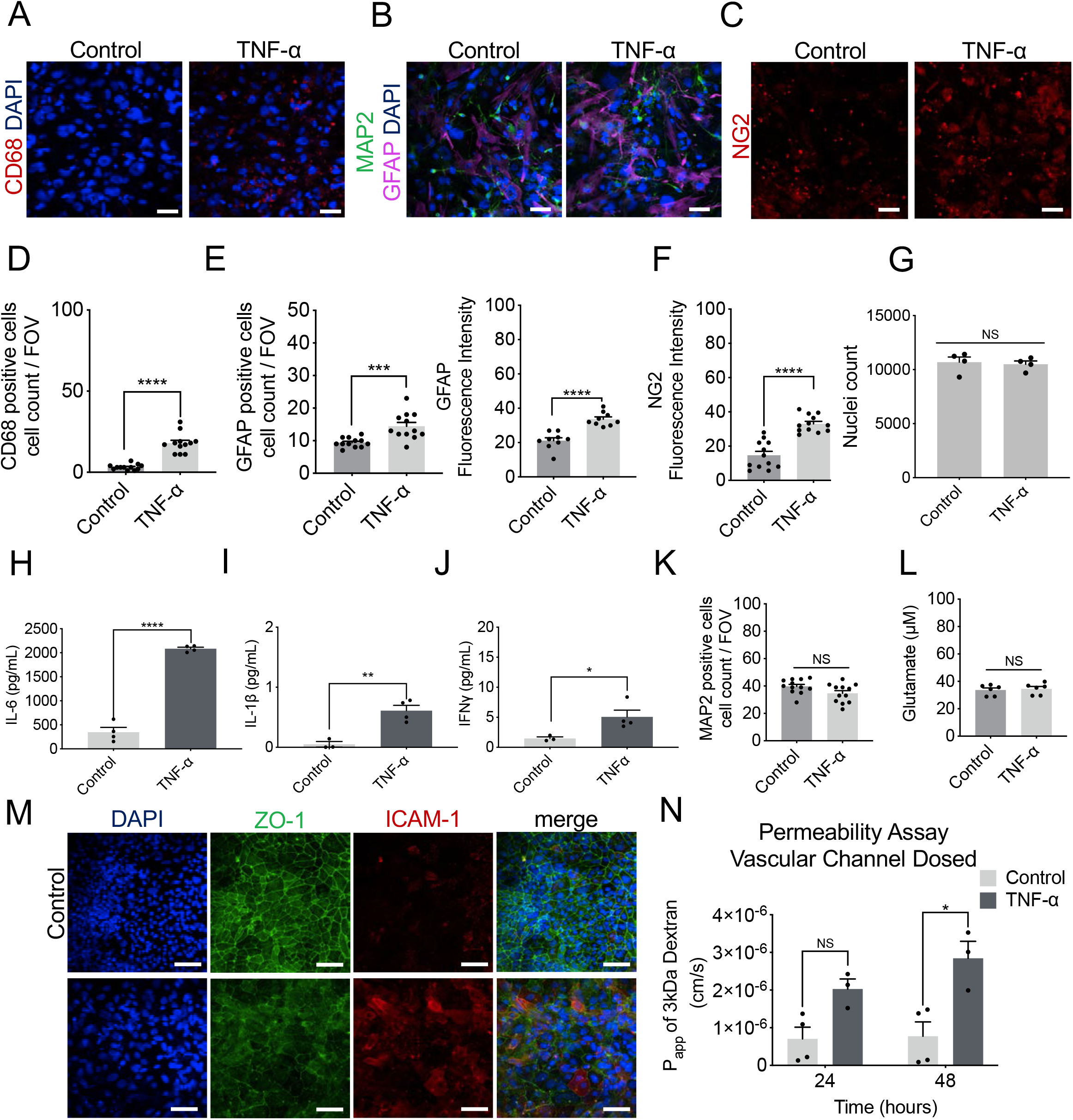
Brain-Chip response to TNF-α perfused through the vascular channel. (**A to C)** Representative immunofluorescent staining for microglia (CD68, red), neurons (MAP2, green), astrocytes (GFAP, magenta), nuclei (DAPI, blue), and pericytes (NG2, red) in TNF-α-treated of control chips (bar, 100 nm). **(D)** Quantification of the CD68-positive events/field of view in n=4 randomly selected different areas/chip, n=3 Brain-Chips; data are expressed as mean ± S.E.M., ****P<0.0001 compared to the untreated control group, statistical analysis by Student’s t-test. **(E)** (Left) Quantification of the number of GFAP-positive events/field of view in n=4 randomly selected different areas/chip, n=3 Brain-Chips; data are expressed as mean ± S.E.M., \*\*\**P*<0.001 compared to the untreated control group; statistical analysis with Student’s t-test. (Right) Quantification of GFAP fluorescent intensity in n=3 randomly selected different areas/chip, n=3 Brain-Chips, \*\*\*\**P*<0.0001 compared to the untreated control group, statistical analysis with Student’s t-test. **(F)** Quantification of fluorescent intensity of NG2 in, n=4, randomly selected different areas/chip, n=3 Brain-Chips; data are expressed as mean ± S.E.M., \*\*\*\**P*<0.0001 compared to the untreated control group; statistical analysis with Student’s t-test. **(G)** The nuclei counts based on DAPI staining were similar between the control and treated groups (n=4 Brain-Chips, data are expressed as mean ± S.E.M., NS: Not Significant compared to the untreated control group). Statistical analysis with Student’s t-test. (**H to J**) Secreted levels of the proinflammatory cytokines IL-6, IL-1β and IFNγ, in the brain channel of control or TNF-α treated Brain-Chips. n=3-4 independent chips, data are expressed as mean ± S.E.M., *P<0.05, P**<0.01, ****P<0.0001, statistical analysis with Student’s t-test. **(K)** Quantification of the number of MAP2 events/field of view in n=4 randomly selected different areas/chip, n=3 Brain-Chips, data are expressed as mean ± S.E.M., NS: Not Significant compared to the untreated control group, statistical analysis with Student’s t-test. **(L)** Levels of secreted glutamate in the brain channel on culture day 7 (n=6 independent chips; data are expressed as mean ± S.E.M., NS: Not Significant compared to the untreated control group). Statistical analysis with Student’s t-test. **(M)** Immunofluorescent staining of cell nuclei (DAPI, blue), Intercellular Adhesion Molecule 1 (ICAM-1, red), tight junction protein 1 (ZO-1, green), and a merged image of all three markers (bar, 100 nm). **(N)** Quantitative barrier function analysis via apparent permeability to 3kDa fluorescent dextran, upon 48h of exposure to TNF-α via the vascular channel. n=3∼4 independent chips, NS= Not Significant, \**P*<0.05, control group compared to TNF-α treated group after 24h and 48 h of treatment; data are expressed as mean ± S.E.M, statistical analysis by two-way ANOVA with Tukey’s multiple comparisons test.

Next, we characterized markers indicative of neuroinflammation. We found significantly higher levels of IFNγ, IL-1β, and IL-6 in the effluent collected from the brain channel, following two days of vascular exposure to TNF-α, compared to the untreated group (**Figure 6H to 6J)**. These findings are consistent with published studies showing that TNF-α crossing through the BBB activates the microglia and induced the release of proinflammatory cytokines, resulting in further propagation of the inflammatory process in the brain (Qin et al., 2007; Tangpong et al., 2006). We also found higher levels of IFNγ, IL-1β, and IL-6 in the vascular channel media in the TNF-treated model group compared to the untreated group, all of which contribute to increase the barrier permeability (**Figure S4B**). However, no neuronal damage was detectable, as per the number of MAP2-positive neurons and the secreted glutamate levels (**Figures 6K and 6L**). Thus, vascular-mediated challenge of the brain with TNF-α induces neuroinflammation, although relatively milder compared to that following direct exposure to TNF-α, administrated through the brain channel.

Immunofluorescence analysis showed significantly attenuated expression of the tight junction protein ZO-1 and increased expression of ICAM-1 in the TNF-α treated chips compared to the control chips (**Figures 6M**). Further, the barrier permeability to 3kDa dextran was significantly increased in the TNF-α treated chips (**Figure 6N**). Overall, these findings were similar to those described in detail above, in the neuroinflammation model above (**Figure 5A, 5B and 5C**).

Brain barriers are uniquely positioned to communicate signals between the central nervous system and peripheral organs. The BBB cells (endothelial cells, pericytes and astrocytes) respond to signals originating from either side by changes in permeability, transport, and secretory functions (Cunningham et al., 2009; Erickson and Banks, 2018; Verma et al., 2006). It has been previously shown that TNF-α crosses the intact BBB by a receptor-mediated transport system, upregulated by CNS trauma and inflammation (Gutierrez et al., 1993; Pan and Kastin, 2002; Pan et al., 2003a). Free traffic of radioactively labelled TNF-α from blood to brain and cerebrospinal fluid (CSF) has been shown in mice (Gutierrez et al., 1993; Pan et al., 1997), as well as in monolayers of cultured cerebral microvessel endothelial cells (Pan et al., 2003b). We measured the levels of TNF-α in effluent from both the brain and vascular channels (**Figure S4C**) and identified significantly lower basal levels in the former (control condition). Our results confirm that TNF-α can cross through the intact barrier in either direction, i.e., brain or vascular. Also, barrier disruption resulted in significantly increased TNF-α levels from the vascular compartment into the brain compartment and vice versa (**Figure 7A and 7B**).

**Figure 7.**
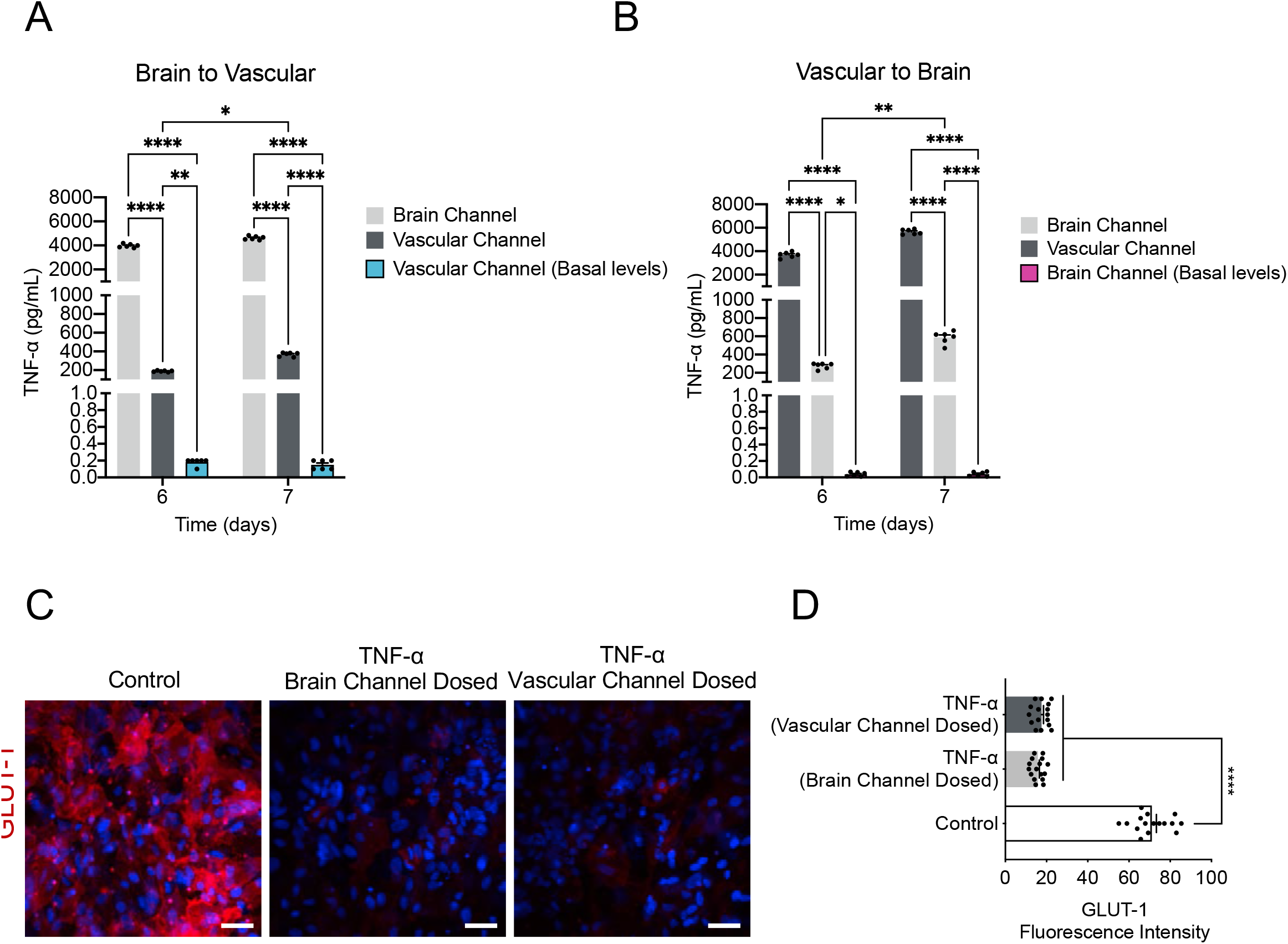
Impact of inflammation on barrier transport systems. (**A, B**) The transport of TNF-α from the brain to vascular or vice versa was shown by assessment of the levels of TNF-α secreted in the vascular or brain channel of control (basal levels) or TNF-α dosed chips either through the brain (brain to vascular) or the vascular channel (vascular to brain), n=6 independent chips, data are expressed as mean ± S.E.M., *P*\*<0.05, *P*\*\*<0.01, \*\*\*\**P*<0.0001, statistical analysis by two-way ANOVA with Tukey’s multiple comparisons test. **(C)** Representative immunofluorescence images of GLUT-1 transporter (red) expression and DAPI (blue), in the endothelial cells of the vascular channel of the Brain Chip upon challenge with TNF-α through the brain or the vascular channels. Vehicle-treated chips serve as control (bar, 100 nm). **(D)** Quantification of the GLUT-1-fluorescence intensity /field of view in 4 randomly selected different areas/chip, n=4 Brain-Chips; data are expressed as mean ± S.E.M., ****P<0.0001 compared to the untreated control group, statistical analysis by two-way ANOVA with Tukey’s multiple comparisons test.

Still, a knowledge gap remains in our in-depth understanding of the interplay between gain/loss of function of drug transporters and their role in the development of neurological diseases, whether changes in transporter expression are a cause or consequence of these diseases. Glucose transporter-1, GLUT-1 is the major cerebral glucose transporter, and is expressed at particularly high levels in endothelial cells that line the brain capillaries (Vannucci, 1994). Although decreases in GLUT-1 expression is a well-accepted biomarker for degenerative and inflammatory brain diseases (Winkler et al, 2015), howit is associated to their pathogenesis remains elusive. To this purpose, we measured the GLUT-1 expression in both TNF-α treatment conditions (brain or vascular side dosing) compared to the control group. Analysis of immunofluorescence images showed substantial decrease in GLUT-1 expression upon TNF-α treatment as compared to the control group (**Figure 7C and 7D**), showing the sensitivity of the Brain-Chip to capture the direct effect of TNF-α on GLUT-1 transporter. Although there are differences associated with the specific route of administration of TNF-α in the inflammation-induced mechanisms, exposure through the brain or the vascular channel represent relevant *in vivo* conditions. All the above taken together, suggest that the Brain-Chip may advance our understanding on the molecular mechanisms underlying function of BBB transporters in basal and disease states.

## DISCUSSION

In the present study, we designed a microfluidic model, a human Brain-Chip, that recreates several functional features of the neurovascular unit. Cortical neurons, astrocytes, microglia, and pericytes compose the parenchymal basement membrane (brain side) whereas astrocytic end-feed embracing the abluminal aspect of the brain microvessels (vascular side). Astrocytes and microglia were maintained in a resting state in coordination with low levels of cytokine secretion. The endothelial-like monolayer within the human Brain-Chip sustained the expression of tight junction proteins and showed low barrier permeability levels for seven days in culture, similar to those reported for the human brain *in vivo* (Shi et al., 2014; Yuan et al., 2009), for seven days in culture.

By leveraging next-generation sequencing data and information retrieved from well-curated databases providing signature gene sets characteristic for the human cortex, we were able to show that the Brain-Chip exhibits higher transcriptomic similarity to the adult cortical tissue than the transwells, both models with the same cellular composition. These data complement previous reports on the advantages of the microfluidic organ-chip systems to provide a better tissue-relevant microenvironment compared to other commonly used conventional culture systems (Pediaditakis et al., 2021; Sances et al., 2018). Most importantly, our findings demonstrate the closeness of the Brain-Chip to the adult human cortex tissue and the cells’ maturation state after seven days in culture.

Neuroinflammation emerges as an essential process in the pathogenesis of neurodegenerative diseases. Several studies have shown direct effects of the activated microglia, astrocytes, and pericytes and the secreted cytokines in the brain and BBB functions (Biswas et al., 2020; Freitas-Andrade et al., 2020; Horng et al., 2017; Liebner et al., 2018; Sofroniew, 2015; Sweeney et al., 2019). TNF-α, a key mediator of inflammation, impairs neuronal function, suppresses long-term hippocampal potentiation (LTP), a mechanism essential for memory storage and consolidation (Cunningham et al., 1996), and affects synaptic transmission (Singh et al., 2019). Further, TNF-α levels have been found markedly elevated in the brains of patients with Alzheimer’s disease (Heneka and O’Banion, 2007), indicative of the active inflammatory process in the disease. It has been proposed that systemic inflammation exacerbates neuroinflammation and neurodegeneration via circulating pro-inflammatory factors (Perry et al., 2007a), such as TNF-α, crossing the BBB via active transport (Osburg et al., 2002; Pan and Kastin, 2007) or through the compromised barrier (Franzén et al., 2003; Trickler et al., 2005). Despite the increasing experimental and clinical evidence on the connection between neuroinflammation, neurodegeneration, and, ultimately, neuronal death, the development of effective therapeutic targets is still slow. A major factor contributing to the latter is that there is still lack of specific models for the onset and progress of human brain diseases. Most of the existing *in vitro* BBB models do not incorporate neurons and glia in the blood-brain barrier cell systems, resulting in incomplete modeling of the inflammatory responses.

In the present study, we characterized the Brain-Chip responses upon exposure to TNF-α via two distinct routes, either directly through the brain channel or via the vascular channel, where TNF-α reaches the brain cells by crossing through the barrier either actively or paracellularly. We show that the brain’s exposure to TNF-α, either directly or through the BBB, results in activation of microglia and astrocytes, secretion of cytokines, and neuronal damage. As expected, exposure to TNF-α induced significant changes in tight junction formation that compromised the barrier permeability and induced adhesion molecules such as ICAM-1, which propagated the inflammatory response by facilitating the recruitment of the immune cells to the brain (Marchetti and Engelhardt, 2020). We expect that future studies set to characterize the precise sequence of events following exposure to systemic- or tissue-induced inflammatory injury might provide important, targetable hints for critical cell-driven mechanisms in neuroinflammation.

The comprehensive Brain-Chip model presented here can enhance our current capability to interrogate both brain barrier dysfunction and neuron-glia interactions underlying the onset and progress of neuroinflammation, for the benefit of human patients.

## METHODS

### Brain-Chip Microfabrication and Zoë^®^ Culture Module

The design and fabrication of Organ-Chips used to develop the Brain-Chip was based on previously described protocols (Huh et al., 2013). The chip is made of transparent, flexible polydimethylsiloxane (PDMS), an elastomeric polymer. The chip contains two parallel microchannels (a 1 × 1 mm brain channel and a 1 × 0.2 mm vascular channel) that are separated by a thin (50 μm), porous membrane (7 μm diameter pores with 40 μm spacing) coated with E.C.M. (400 µg/mL collagen IV, 100 µg/mL fibronectin, and 20 µg/mL laminin, at the brain and vascular side). Brain-Chips were seeded with human iPSC-derived glutamatergic and GABAergic neurons (NeuCyte;1010) at a density of 4×10^6^ cells/mL and 2×10^6^ cells/mL respectively, co-cultured with, human primary astrocytes (NeuCyte;1010) at a density of 2×10^6^ cells/mL, human microglial cell line (ATCC; CRL3304) at a density of 2×10^5^ cells/mL, and primary pericytes (Sciencell;1200) at a density of 1.5×10^5^ cells/mL, using “seeding medium” (NeuCyte), and incubated overnight. The next day, human iPSC-derived Brain Microvascular Endothelial-like cells were seeded in the vascular channel at a density of 14 to 16×10^6^ cells/mL using human serum-free endothelial cell medium supplemented with 5% human serum from platelet-poor human plasma (Sigma) and allowed to attach to the membrane overnight. Chips were then connected to the Zoë^®^ Culture Module (Emulate Inc.). At this time, the medium supplying the brain channel was switched to maintenance medium (Neucyte), and the serum of the vascular medium was lowered to 2%. Chips were maintained under constant perfusion at 60 µL/h through both chips’ brain and vascular channels until day seven.

### Brain Transwell Model

The conventional cell cultures (transwells) and the Brain-Chips were seeded using the same ECM composition as well as cell composition, media formulations and seeding density. At the first experimental day (D0) the cortical (Glutamatergic and GABAergic subtypes) neurons, astrocytes, microglia, and pericytes were seeded on the apical side, followed by the seeding of the endothelial cells (D1) on the basolateral side of the 0.47 cm2 Transwell-Clear permeable inserts (0.4-μm pore size). For the apical compartment we used NeuCyte medium, while for the basolateral compartment we used hESFM with 5% human serum from platelet-poor human plasma. The cells maintained under static conditions throughout the duration of the experiment (D8). The culture medium was replaced daily in both compartments.

### Differentiation of iPSCs into Brain Microvascular Endothelial-like Cells

Human iPSCs (Donor 1: RUCDR; ND50028, Donor 2: iXcell; 30HU-002) were passaged onto Matrigel in mTeSR1 medium for 2 to 3 days of expansion. Colonies were singularized using Accutase (STEMCELL; 07920) and replated onto Matrigel-coated plates at a density 25-50 × 103 cells/cm2 in mTeSR1 supplemented with 10 mM Rho-associated protein kinase (ROCK) inhibitor Y-27632 (STEMCELL; 72304). Singularized Human iPSCs were expanded in mTeSR1 for 3 days. Cells were then treated with 6 mM CHIR99021 (STEMCELL; 72052) in DeSR1: DMEM/Ham’s F12 (Thermo Fisher Scientific; 11039021), 1X MEM-NEAA (Thermo Fisher Scientific; 10370021), 0.5% GlutaMAX (Thermo Fisher Scientific; 35050061), and 0.1 mM b-mercaptoethanol (Sigma). On Day 1, the medium was changed to DeSR2: DeSR1 plus 1X B27 (Thermo Fisher Scientific) daily for another 5 days. On day 6, the medium was switched to hECSR1: hESFM (ThermoFisher Scientific) supplemented with bFGF (20 ng/mL), 10 mM Retinoic Acid, and 1X B27. On day 8, the medium was changed to hECSR2 (hECSR1 without R.A. or bFGF). On day 10 cells were dissociated with TrypLE(tm) and plated at 1×10^6^ cells/cm2 in hESFM supplemented with 5% human serum from platelet-poor human plasma onto a mixture of collagen IV (400µg/mL), fibronectin (100µg/mL), and laminin (20 µg/mL) coated flasks at a density of 1×10^6^ cells/cm2. After 20 min the flasks were rinsed using hESFM with 5% human serum from platelet-poor human plasma with Y-27632 as a selection step to remove any undifferentiated cells and allowed to attach overnight (Qian et al., 2017).

### Morphological Analysis

Immunocytochemistry was conducted as previously described (Pediaditakis et al., 2021). Cells were blocked on the Brain-Chip in phosphate-buffered saline (PBS) containing 10% donkey serum (Sigma) at 4°C overnight. Saponin 1% was used to permeabilize membrane when required. Primary antibodies were MAP2 (Thermo Fisher Scientific; MA512826), VGLUT1 (Thermo Fisher Scientific; 48-2400), Synaptophysin (Abcam; 32127), GFAP (Abcam; ab53554), GLAST (Invitrogen; PA5-19709), s100β (Abcam; 52642), NG2 (Abcam; ab83178), αSMA (Abcam; 7817), IBA1 (FUJIFILM; 019-19741), CD68 (Abcam; ab213363), ICAM-1 (R&D Systems; BBA3), Ki67 (Abcam; 197234), ZO-1 (Thermo Fisher Scientific; 402200), Occludin (Invitrogen; OC-3F10), Claudin-4 (Invitrogen; 329494), TRPV6 (Proteintech;13411-1-AP), PECAM1 (Thermo Fisher Scientific; RB-1033-P1), CD11b (Invitrogen; MA1-80091), CD45 (Invitrogen; 17-0409-42), GLUT1 (Thermo Fisher Scientific; SPM498), P-gp (Thermo Fisher Scientific; p170(F4)), MRP-1 (Millipore; MAB4100), Transferrin receptor (Abcam; 216665). Chips treated with corresponding Alexa Fluor secondary antibodies (Abcam) were incubated in the dark for 2 h at room temperature. Cells were then counterstained with nuclear dye DAPI. Images were acquired with an inverted laser-scanning confocal microscope (Zeiss LSM 880).

### Flow Cytometry

Cells were dissociated with Accutase, fixed in 1% PFA for 15 min at room temperature, and then washed with 0.5% bovine serum albumin (BSA) (Bio-Rad) plus 0.1% Triton X-100 three times. Cells were stained with primary and secondary antibodies diluted in 0.5% BSA plus 0.1% Triton X-100. Data were collected on a FACSCelesta flow cytometer (Becton Dickinson) and analyzed using FlowJo. Corresponding isotype antibodies were used as FACS (fluorescence-activated cell sorting) gating control. Details about antibody source and usage are provided in table.

### Visualization of Transferrin Receptor Internalization

Human iPSC-derived Brain Microvascular Endothelial-like cells were treated with 25 μg/mL fluorescent transferrin conjugate (Thermo Fisher Scientific) and incubated at 37oC for 30 min. Cells were washed twice with LCIS and fixed with P.F.A. Cells labeled with Alexa Fluor(tm) Plus 647 Phalloidin and DAPI and then imaged with Zeiss LSM 880.

### Scanning Electron Microscopy

At the indicated timepoints Brain-Chips were fixed at room temperature, for 2 hours in 2.5% Glutaraldehyde solution and washed three times with 0.1M sodium cacodylate (NaC) buffer. Concomitantly, the chip was trimmed using a razor so that the lateral and top chunks of PDMS are removed and the top channel is revealed. Afterwards, the samples were fixed with 1% osmium tetroxide (OsO4) in 0.1M NaC buffer for 1 hour at room temperature and dehydrated in graded ethanol. The chip samples were dried using the chemical drying agent Hexamethyldisilizane (HMDS), sputter coated with platinum and images were acquired using the Hitachi S-4700 Field Emission Scanning Electron Microscope.

### Permeability Assays

To evaluate the establishment and integrity of the barrier, 3 kDa Dextran, Cascade Blue, was added to the vascular compartment of the Brain-Chip at 0.1 mg/mL. After 24 h, effluent from both channels was sampled to determine the dye’s concentration that had diffused through the membrane. The apparent paracellular permeability (Papp) was calculated based on a standard curve and using the following formula:

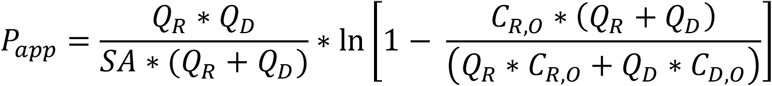

where SA is the surface area of sections of the channels that overlap (0.17cm2), Q_R_ and Q_D_ are the fluid flow rates in the dosing and receiving channels respectively, in units of cm3/s, C_R,O_ and C_D,O_ are the recovered concentrations in the dosing and receiving channels respectively, in any consistent units.

### TNF-α Treatment

To mimic the inflammatory condition, cells were treated on either brain or vascular channel with TNF-α (Tumor Necrosis Factor-α, R&D Systems; 210-TA). The treatment was initiated after the formation of a confluent monolayer at ∼5 days in culture. Cells were further incubated in a culturing medium, including TNF-α (100 ng/mL) up to 48 h.

### Western Blotting

RIPA cell lysis buffer supplemented with protease and phosphatase inhibitors (Sigma) was used for the extraction of total protein from either brain or vascular channel. The Auto Western Testing Service was provided by RayBiotech, Inc. (Peachtree Corners, GA USA). 0.2 mg/mL sample concentration was loaded into the automated capillary electrophoresis machine. Glial fibrillary acidic protein (GFAP) and Gluceraldehyde-3-phosphate dehydrogenase (GAPDH) antibody provided by RayBiotech was used as the loading control.

### ELISA Analysis

The levels of IFNγ, IL-1β, and IL-6 were measured by M.S.D. 96-well plate Human Pro-Inflammatory V-PLEX Human Pro-Inflammatory Assay kits. The secreted levels of Glutamate were measured by Glutamate Assay Kit (Fluorometric) (Abcam; ab138883).

### RNA Isolation and Sequencing

According to manufacturer’s guidelines, we used TRIzol (TRI reagent, Sigma) to extract the RNA. The collected samples were submitted to GENEWIZ South Plainfield, NJ, for next-generation sequencing. After quality control and RNA-seq library preparation the samples were sequenced with Illumina HiSeq 2×150 system using sequencing depth ∼50M paired-end reads/sample.

### RNA Sequencing Bioinformatics

Using Trimmomatic v.0.36 we trimmed the sequence reads and filtered-out all poor-quality nucleotides and possible adapter sequences. The remained trimmed reads were mapped to the Homo sapience reference genome GRCh38 using the STAR aligner v2.5.2b. Next, using the generated BAM files we calculated for each sample the unique gene hit-counts by using the featureCounts from the Subread package v.1.5.2. It is worth noting that only unique reads that fell within the exon region were counted. Finally, the generated hit-counts were used to perform DGE analysis using the “DESeq2” R package (Love et al., 2014). The thresholds used for all the DGE analyses were: |log_2_(Fold Change)| ≥ 1 and adjusted p-value ≤ 0.01.

### GO term Enrichment Analysis

The DE genes identified after performing the DGE analyses were subjected to Gene Ontology (GO) enrichment analysis. The GO terms enrichment analysis was performed using the Gene Ontology knowledgebase (Gene Ontology Resource http://geneontology.org/).

### GTEx Portal provides 255 RNA-seq Samples for Human Brain-Cortex

From these samples only 5 were from healthy individuals and had RNA Integrity Number larger than 8 (RIN ≥ 8), which indicates good RNA-quality. To create a “balanced” dataset (i.e., 4 samples per condition) from these 5 samples, we selected the group of 4 that had the smallest variance. We combined the selected samples with the 16 samples from our healthy models (i.e., Brain-Chips and transwells on Days 5 and 7). Next, we used the “remove Batch Effect” function of the “limma” R package (Ritchie et al., 2015) to remove shifts in the means between our samples (Brain-Chips and transwells) and the 4 human Brain-Cortex samples retrieved from GTEx portal (Lonsdale et al., 2013). The same process was repeated to combine the 16 samples from our healthy models with each one of the two different cerebral cortex RNA-seq data from two different human donors (donor 9861 and 10021) available in the Allen Brain Atlas (© 2010 Allen Institute for Brain Science. Allen Human Brain Atlas. Available from: human.brain-map.org). This dataset used for the calculation of the Transcriptomic Signature Distances (TSDs) (Manatakis et al., 2020).

### Transcriptomic Signature Distance (TSD) Computation

TSD is a novel distance metric based on information theory that allows us to reliably assess the transcriptomic similarity between organ tissue samples. The TSD uses (i) next-generation sequencing data and (ii) tissue-specific genes (i.e., signature genes) provided by the well-curated and widely accepted Human Protein Atlas (HPA) project (Uhlén et al., 2015), and calculates the transcriptomic distance of a tissue sample (e.g., Brain-Chip or transwell) from the reference tissue (in our case the Human Brain-Cortex). As signature genes we used the set of 2587 genes that are reported to have significantly elevated expression levels in the brain tissue compared to other tissue types (Uhlén et al., 2015).

### Statistical Analysis

All experiments were performed in triplicates. Analysis of significance was performed by using two-way ANOVA with Tukey’s multiple comparisons test or unpaired t-test depending on the data sets. The error bars represent standard error of the mean (s.e.m); p-values < 0.05 and above were considered significant.

## Data Availability

All data generated or analyzed during this study are included in this published article. RNA sequencing data have been deposited in the National Center for Biotechnology Information Gene Expression Omnibus (GEO).

## Code Availability

All the code for the analysis in this report is derived from previously published reports. It is also explained and cited in the appropriate material and methods section.

## AUTHOR CONTRIBUTION

I.P. developed the Brain-Chip model, designed, and performed experiments, collected, and analyzed data, and wrote the paper. K.R.K. contributed to the Brain-Chip model development, performed experiments, collected data, and contributed to the writing of the paper. D.V.M. processed and analyzed the transcriptomic data, incorporated the associated data in the manuscript and contributed to the writing of the paper. C.Y.L, S.B. and A.S. helped perform experiments. E.S.M. was involved in the bioinformatic analysis and contributed to the writing of the paper. A.G., L.E., L.L.R provided critical feedback and reviewed the manuscript. C.D.H. provided insightful input on the engineering aspects of the project. K.K. supervised the project, contributed to the Brain-Chip model development, and wrote the paper.

## ACKNOWLEDGMENTS

We would like to thank Dr. Athanasia Apostolou for providing technical assistance with the scanning electron microscope. This work was supported by the National Institute of Health, National Center for Advancing Translational Sciences (UG3TR002188; to C.D.H. and K.K.). The content is solely the responsibility of the authors and does not necessarily represent the official views of the National Institutes of Health. We also thank Brett Clair for scientific illustrations.

