## Supplementary Information for "A Microengineered Brain-Chip to Model Neuroinflammation in Humans"

#### SUPPLEMENTAL INFORMATION

**Supplementary Figure. 1. Characterization of endothelial cells and barrier properties**

(**A**) Top: Representative merged confocal image of the brain endothelial-like cells on chip, stained with epithelial markers CADH1 (magenta), TRPV6, Claudin-4 and DAPI (blue) compared to bottom: Human epithelial cells stained positive for the same markers (bar,100 µm).

(**B**) Assessment of the permeability of the Brain-Chip on culture day 7, in the absence or presence of microglia, astrocytes or neurons; n=4-8 independent chips; data are expressed as mean ± S.E.M., ****P<0.0001, NS: Not Significant compared to full model (Brain-Chip), statistical analysis by one-way ANOVA with Sidak's multiple comparisons test.

(**C**) Comparison of the permeability of two different iPSC-derived microvascular endothelial-like cells (iBMECs) culture models, monoculture versus culture in the Brain-Chip (n=4-6 independent chips, ****$P$<0.0001). Data are expressed as mean ± S.E.M, statistical analysis with Student's t-test.

(**D**) Representative merged confocal image of the vascular channel on culture day 7, stained for the tight junction protein marker (ZO-1, green) and Glucose transporter (GLUT-1, red) (bar, 100 µm).

(**E**) Representative merged confocal image of immunofluorescent staining for transferrin (transferrin conjugate, pHrodo, (red) in the cytoplasm of iBMECs, together with F-actin (green) and DAPI (blue) (bar, 100 µm).

(**F)** Flow cytometry analysis of the human brain endothelial-like cells labeled with antibodies for receptors and transporters, following culture on-chip for 7 days.

**Supplementary Figure 2. Comparative analyses of the Transcriptomic profiles of the Brain-Chip, adult cortex tissue and transwell culture.**

(**A**) Boxplots summarizing the distributions of the corresponding pairwise TSD distances. In each pair, one sample belongs to the reference tissue (Human Brain-Cortex; Allen Brain Atlas) and the other either to the reference tissue or to one of our culture models, i.e., Brain-Chip or transwell, from culture days 5 and 7. The Brain-Chip and transwell cultures were run in parallel ****P<0.0001.

**Supplementary Figure 3. Brain-Chip response to TNF-α perfused through the brain channel**

(**A**) Schematic of the timeline of a typical inflammation experiment.

(**B**) Quantification of fluorescent intensity of GFAP in n=3 randomly selected different areas/chip, n=3 Brain-Chips; data are expressed as mean ± S.E.M., ****$P$<0.0001 compared to the untreated control group; statistical analysis with Student's t-test.

(**C**) Western blotting analysis of cell lysates from the brain channel shows increased expression of GFAP levels (MW: 50kDa) following exposure to TNF-α. For loading control, equal amounts of protein were immunoblotted with GAPDH antibody (MW: 37kDa). The numbers 0.26 and 0.69 indicate quantification for relative levels of signal intensity.

(**D**) Nuclei counts as per DAPI staining, are similar in control and TNF-α treated groups (n=4 Brain-Chips; data are expressed as mean ± S.E.M., NS: Not Significant to the untreated control group. Statistical analysis with Student's t-test.
(**E**) Immunocytochemical staining of Ki67-positive cells in control and TNF-α challenged groups (bar, 100 µm). White arrows show Ki67-positive cells.
(**F**) Ki67-positive cells in control and TNF-α challenged Brain-Chips, as percentage of the total number of cells (n=4 Brain-Chips; data are expressed as mean ± S.E.M., NS: Not Significant compared to the control group; statistical analysis with Student's t-test.

**Supplementary Figure 4. Brain-Chip response to TNF-α perfused through the vascular channel**
(**A**) Schematic illustration of perfusion of TNF-α through the vascular channel (BBB) of the Brain-Chip.
(**B**) Secreted levels of proinflammatory cytokines (IL-6, IL-1β, and IFNγ) in the vascular channel of control or TNF-α treated Brain-Chips (either through the brain or vascular channel). n=6 independent chips, data are expressed as mean ± S.E.M., NS: Not Significant, $P$**<0.01, $P$***<0.001, ****$P$<0.0001, statistical analysis with Student's t-test.
(**C**) Quantification of basal TNF-α levels present in the effluent of the Brain or Vascular Channel, on day 5, 6 and 7 of culture. n=3-5 independent chips; data are expressed as mean ± S.E.M. NS: Not Significant, **P<0.01, ***P<0.001, ****P<0.0001; statistical analysis with Student's t-test.

### Supplementary Figure 1

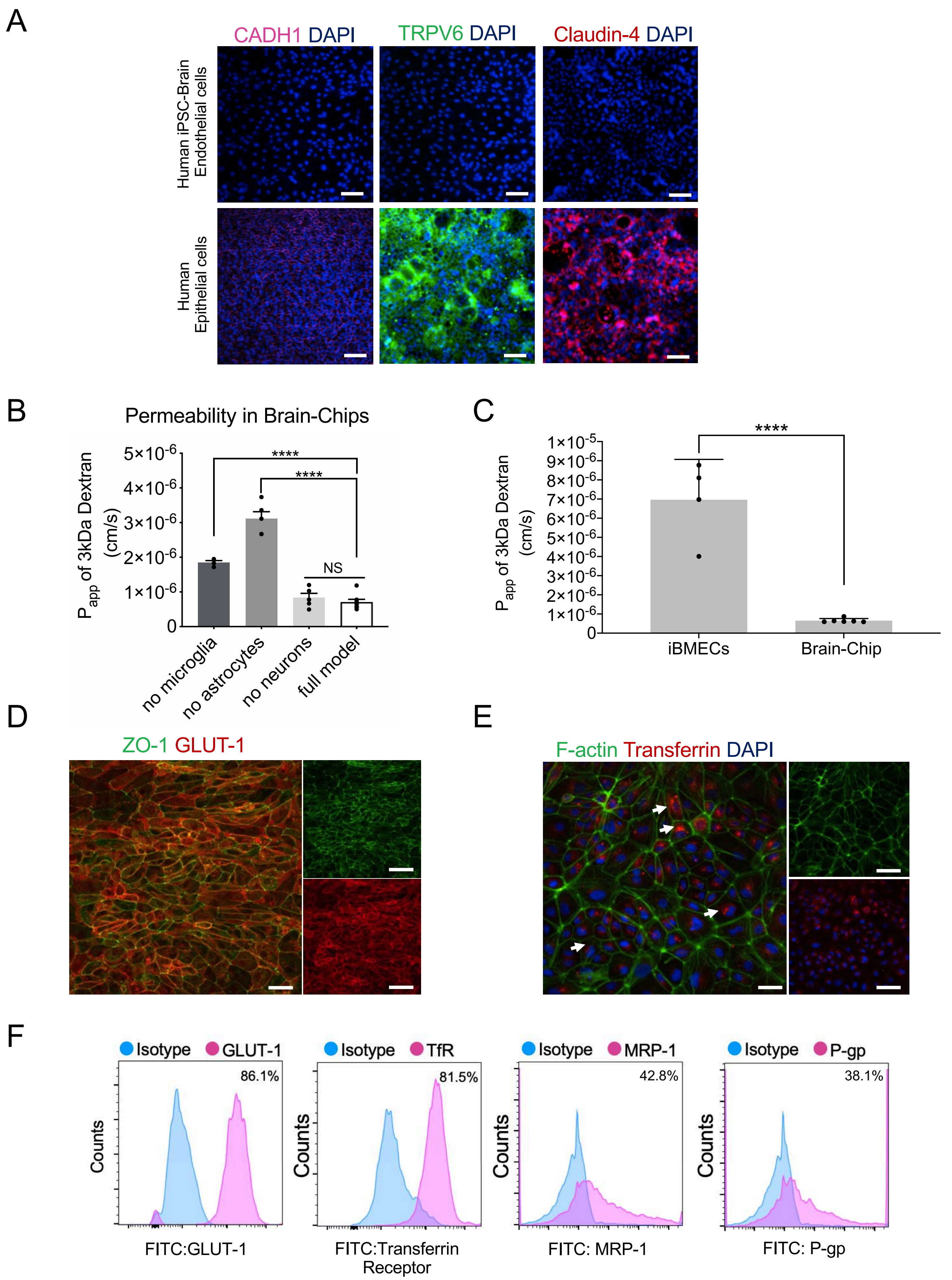

Supplementary Figure 2

A

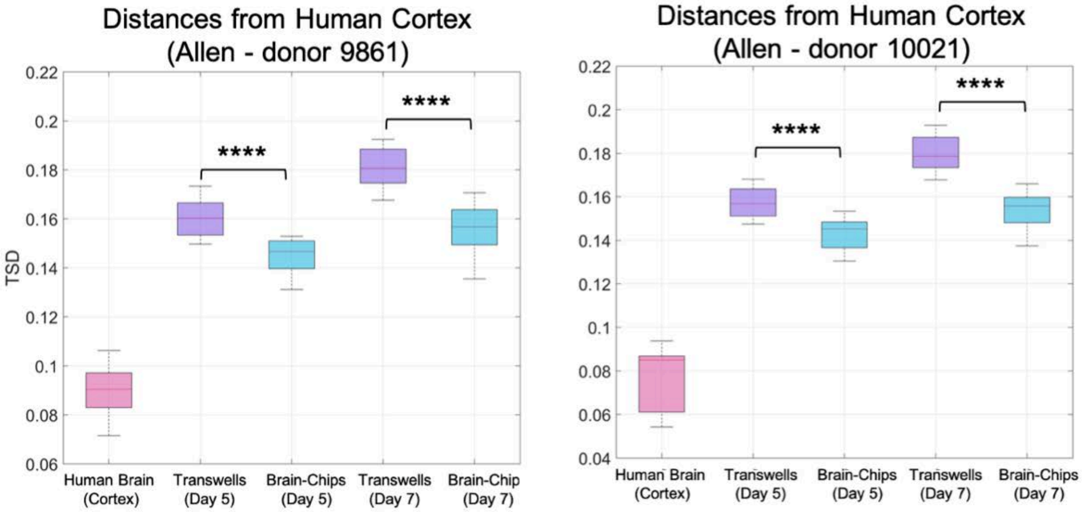

Supplementary Figure 3

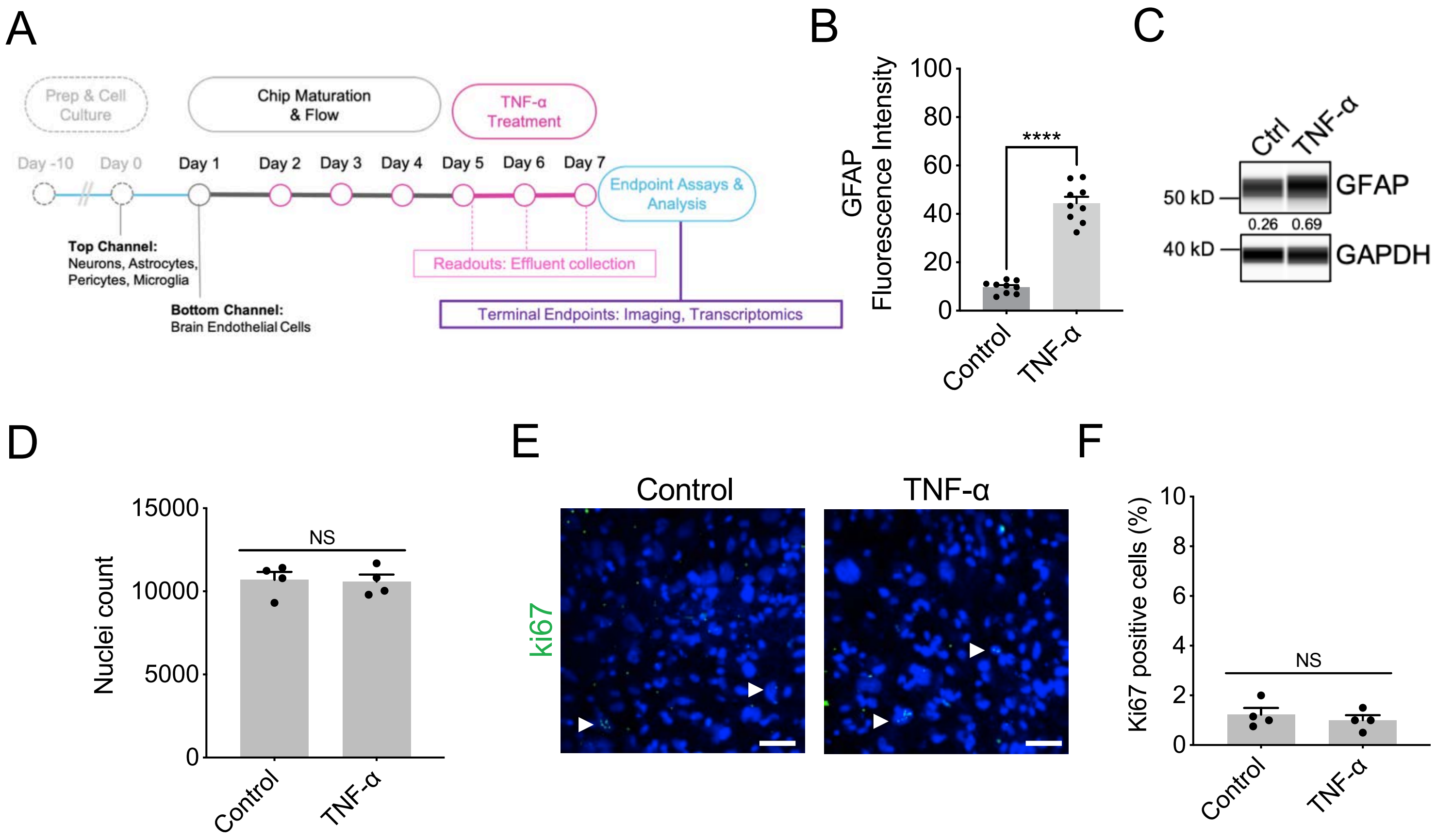

### Supplementary Figure 4

A

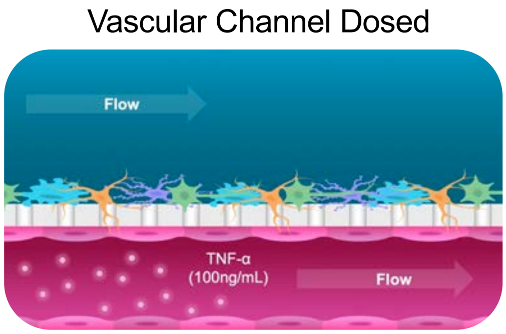


B

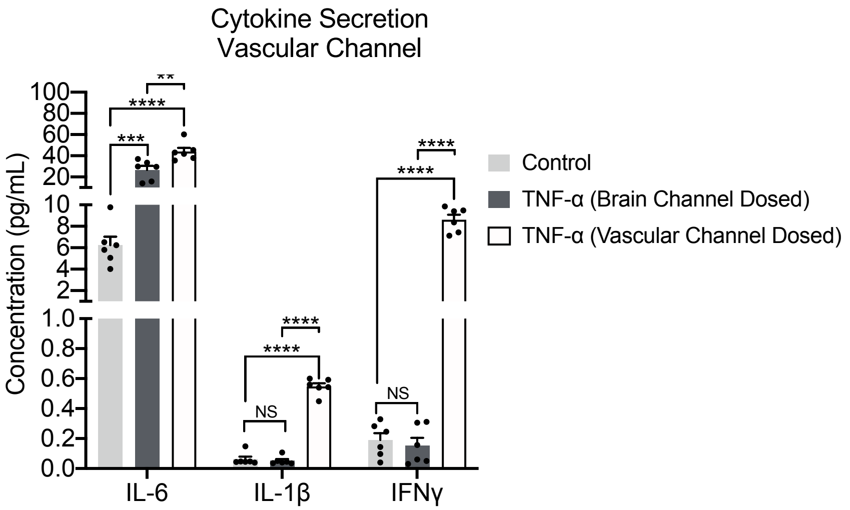


C

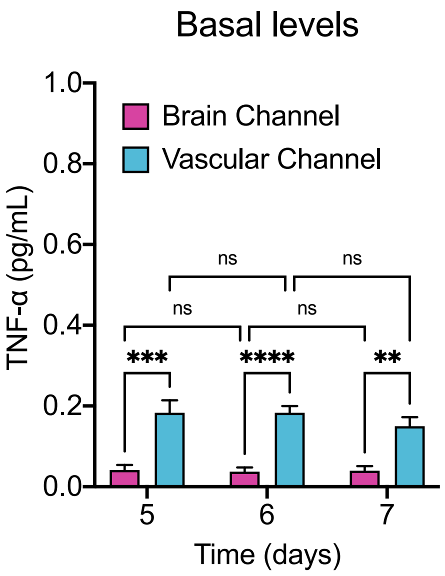
